# ADPriboDB: The Database of ADP-ribosylated Proteins

**DOI:** 10.1101/063768

**Authors:** Christina A. Vivelo, Ricky Wat, Charul Agrawal, Hui Yi Tee, Anthony K. L. Leung

## Abstract

ADP-ribosylation refers to the addition of one or more ADP-ribose units onto proteins post-translationally. This protein modification is often added by ADP-ribosyltransferases, commonly known as PARPs, but it can also be added by other enzymes, including sirtuins or bacterial toxins. While past literature has utilized a variety of methods to identify ADP-ribosylated proteins, recent proteomics studies bring the power of mass spectrometry to determine sites of the modification. To appreciate the diverse roles of ADP-ribosylation across the proteome, we have created ADPriboDB—a database of ADP-ribosylated proteins (http://ADPriboDB.leunglab.org). Each entry of ADPriboDB is annotated manually by at least two independent curators from the literature between January 1975 and July 2015. The current database includes over 12,400 protein entries from 459 publications, identifying 2,389 unique proteins. Here we describe the structure and the current state of ADPriboDB as well as the criteria for entry inclusion. Using this aggregate data, we identified a statistically significant enrichment of ADP-ribosylated proteins in non-membranous RNA granules. To our knowledge, ADPriboDB is the first publicly available database encapsulating ADP-ribosylated proteins identified from the past 40 years, with a hope to facilitate the research of both basic scientists and clinicians to better understand ADP-ribosylation at the molecular level.

## INTRODUCTION

ADP-ribosylation refers to the addition of one or more ADP-ribose units onto proteins through the transfer of the ADP-ribose group from NAD^+^ to target proteins post-translationally. ADP-ribose groups can be attached singly as mono(ADP-ribose) or in polymeric chains as poly(ADP-ribose) (PAR) by the enzymatically active members of the family of 17 human ADP-ribosyltransferases, commonly known as PARPs (1–3). In addition, mono(ADP-ribosyl)ation can be facilitated by other enzymes including sirtuins, extracellular membrane-associated ADP-ribosyltransferases, and bacterial toxins (4–6). Of note, non-enzymatic addition of ADP-ribose groups onto proteins was also reported in vitro (7, 8). ADP-ribosylation can be added onto amino acids of diverse chemistry, including glutamate, aspartate, lysine, arginine and cysteine (9, 10). Some of these ADP-ribose conjugations have shown to be reversible and their removal is mediated by two broad classes of enzymes that cleaves PAR or the bond between ADP-ribose and its conjugated amino acid (11, 12). Poly(ADP-ribose) glycohydrolase (PARG) and ADP-ribosylhydrolase 3 (ARH3) cleave the ribose-ribose bonds between ADP-ribose subunits (11, 12). While ADP-ribosylhydrolase ARH1 removes single ADP-ribose groups from conjugated arginine residues, ADP-riboylhydrolases MacroD1, MacroD2, TARG1 remove the last remaining ADP-ribose groups from poly(ADP-ribosyl)ated substrates or single ADP-ribose groups from mono(ADP-ribosyl)ated proteins conjugated at acidic residues (11, 12). TARG1 is unique for its additional ability to remove the whole PAR chain specifically at glutamate-ADP-ribose ester bonds (13). Several techniques have been developed to enrich and identify ADP-ribosylated substrates (reviewed in (14)). Some techniques use antibodies or protein domains that bind ADP-ribose to pull out the modification from cell lysates and use mass spectrometry to identify ADP-ribosylated substrates (15–17). Other techniques utilize protein microarrays and recombinant PARPs to identify PARP-specific substrates (18, 19).

ADP-ribosylation has numerous and diverse effects on protein functions and cellular pathways, including DNA damage, transcription, chromatin organization, stress responses, circadian rhythms, cell cycle regulation and RNA metabolism (2, 20–24). The addition of the modification can affect the substrate’s stability and activity, as well as serve as a scaffold to recruit other proteins non-covalently (20, 25–27). ADP-ribosylation activity spans a diverse range of organisms across different kingdoms, from viruses to bacteria to mammals, highlighting the importance of this modification in living organisms (28, 29). Because of the diverse roles they play in the cell, ADP-ribosylation and ADP-ribosyltransferase/PARP activity have been implicated in a spectrum of disease pathogenesis, such as cancer, inflammatory diseases and neurological disorders (30, 31). Notably, PARP inhibitors have already shown clinical benefits in multiple cancers, including Olaparib (Lynparza™) which has already been approved by the Food and Drug Administration in the United States and European Medicines Agency (32, 33). Thus, understanding substrate specificity of PARPs and how protein function is affected by ADP-ribosylation are of timely importance and have the potential to improve our understanding of the biology of diseases.

Though the modification was discovered over 50 years ago (34, 35), it was only until recently that several proteomics techniques have been developed to identify the ADP-ribosylation sites (reviewed in (9, 36)). Such site information will provide initial insights to generate hypotheses for testing the function of ADP-ribosylation in modified substrates. To facilitate researchers in appreciating the depth and breadth of ADP-ribosylation’s role across the proteome, we have created ADPriboDB, a database of ADP-ribosylated proteins curated from the literature between 1975 and July 2015 (ADPriboDB.leunglab.org). ADPriboDB is freely accessible online and provides information for each entry on the protein name, gene symbol, UniProt ID, species/cell line of origin, experimental details, and states the original figure/table from which the entry was curated. Entries from mass spectrometry studies include the modified residues and peptide sequenced wherever available. The database curation effort is ongoing and the website also allows single entries or bulk format upload for publication-associated datasets. It is our hope that ADPriboDB will offer insights into the ADP-ribosylated proteome, and may reveal patterns of ADP-ribosylation across substrate proteins, and highlight important interactions and networks, which will aid in our understanding of the myriad roles of ADP-ribosylation played in cells.

## Criteria for inclusion in the ADPriboDB

ADPriboDB was compiled through a manual curation of the literature accessed from PubMed, using the following sixteen search terms: “PARylation” or “PARsylation” or “poly(ADP-ribosyl)ation” or “poly-adp-ribosylation” or “poly(ADP-ribosylation)” or “PARylated” or “PARsylated” or “poly(ADP)ribosylation” or “MARylation” or “MARsylation” or “mono(ADP-ribosyl)ation” or “mono-adp-ribosylation” or “mono(ADP-ribosylation)” or “MARylated” or “MARsylated” or “mono(ADP)ribosylated”. This parameter was further limited for our search between January 1975 and July 2015 yielding 1,456 articles. Each journal article was assessed for the inclusion in ADPriboDB by at least two independent curators.

Diverse techniques have been applied to demonstrate that the entries are ADP-ribosylated (“Method”; see Table S1), including immunoprecipitation with anti-PAR antibodies from cells and autoradiography of *in vitro* modification of recombinant proteins by ADP-ribosyltransferases with ^32^P-labeled NAD^+^. Recent years have seen the application of mass spectrometry approaches for validation of protein modifications and site identification. For the sake of comprehensiveness, ADPriboDB includes all proteins identified from immunoprecipitation/affinity-based approaches as tentatively ADP-ribosylated. The underlying caveat of these approaches using antibodies and ADP-ribose binding domains is that it cannot distinguish whether these proteins are non-covalently bound with ADP-ribose groups or covalently attached to them. However, such distinction can now be made possible with the ability to identify the ADP-ribosylation sites (9, 36). As a result, we include modification details based on mass spectrometry analyses in ADPriboDB whenever they are available. For example, CEBPB was identified in an affinity-based proteomics study using the Af1521 ADP-ribose binding macrodomain (15), but the ADP-ribosylation site was subsequently identified in a global site mapping study (37) (Figure 1).

**Figure 1.**
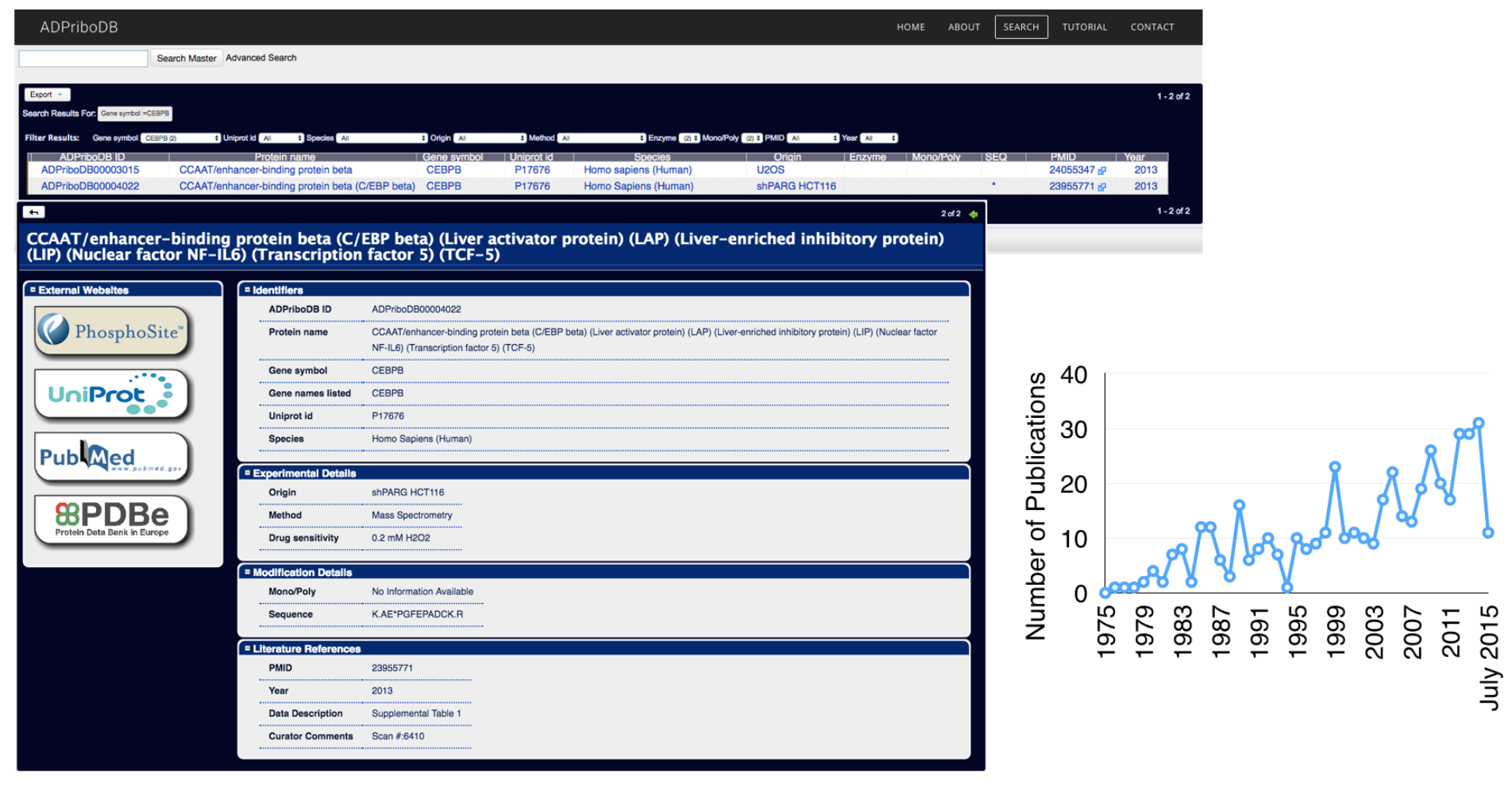
Left: Snapshots of ADPriboDB. Right: The number of publications identifying ADP-ribosylated protein substrates each year from January 1975 through July 2015, totaling to 459 publications.

## ADPriboDB Structure

The database is divided into four sections (Figure 1): “Identifiers”, “Experimental Details”, “Modification Details” and “Literature References”. If a modified protein was identified, the protein name, gene symbol, and corresponding UniProt ID were included in the “Identifiers” section. In the circumstance in which a protein was identified but no UniProt IDs and gene symbols were associated in the publication, the curator(s) determined the best assignment of a UniProt ID based on all the information provided, and noted that this entry as “self-assigned”, along with any other comments in the “Curator Comments” field. If multiple variants of the protein are available on UniProt, the first variant was chosen.

Several key experimental details were collected under the “Experimental Details” section, including the origin of the cell line/tissue (“Origin”) and the species they are derived from (“Species”). Additionally, we noted whether the modification is induced/inhibited by certain drug treatment (“Drug sensitivity”). Under the “Modification Details” section, we noted whether the modification is regulated by certain ADP-ribosyltransferases (“Enzymes”) and whether the modification is mono(ADP-ribosyl)ation or poly(ADP-ribosyl)ation (“Mono/Poly”). In addition, the ADP-ribosylation sites and peptide sequences identified by mass spectrometry were included (“Sequencing Information”). Under the “Literature References” section, the figure/table number displaying the data for modification and an excerpt of the text describing the data were included under “Data Description”. The PubMed ID and year of these publications were included to allow users to easily review or cite the primary sources for the identification of specific ADP-ribosylated proteins.

All entries, given a unique ADPriboDB identification number, were organized in a flat text file database. A user search interface was then created using Xataface, a web application framework for implementing the open-source relational database management system MySQL. ADPriboDB can be searched by Protein Name, Gene Symbol, UniProt ID, PubMed ID and Enzymes (e.g. PARP1). The rest of the webpages to explain the functioning of the database were created through Weebly.

## Current State of ADPriboDB

Manual curation of literature between January 1975 and July 2015 identified 12,428 entries, in which 8,867 of them have protein sequencing data available. The current database entries are derived from 459 publications with experimental supports for PARylated and/or MARylated proteins. The database currently includes 2,389 unique proteins and 573 of them are associated with sequencing information. Though 93% entries are derived from human, over 860 entries are from diverse organisms, which include not only the common model systems, such as mouse, rat, fruitfly and Arabidopsis, but also quail, cow, chicken, sea urchin, octopus, various bacteria and viruses (Figure 2a).

**Figure 2.**
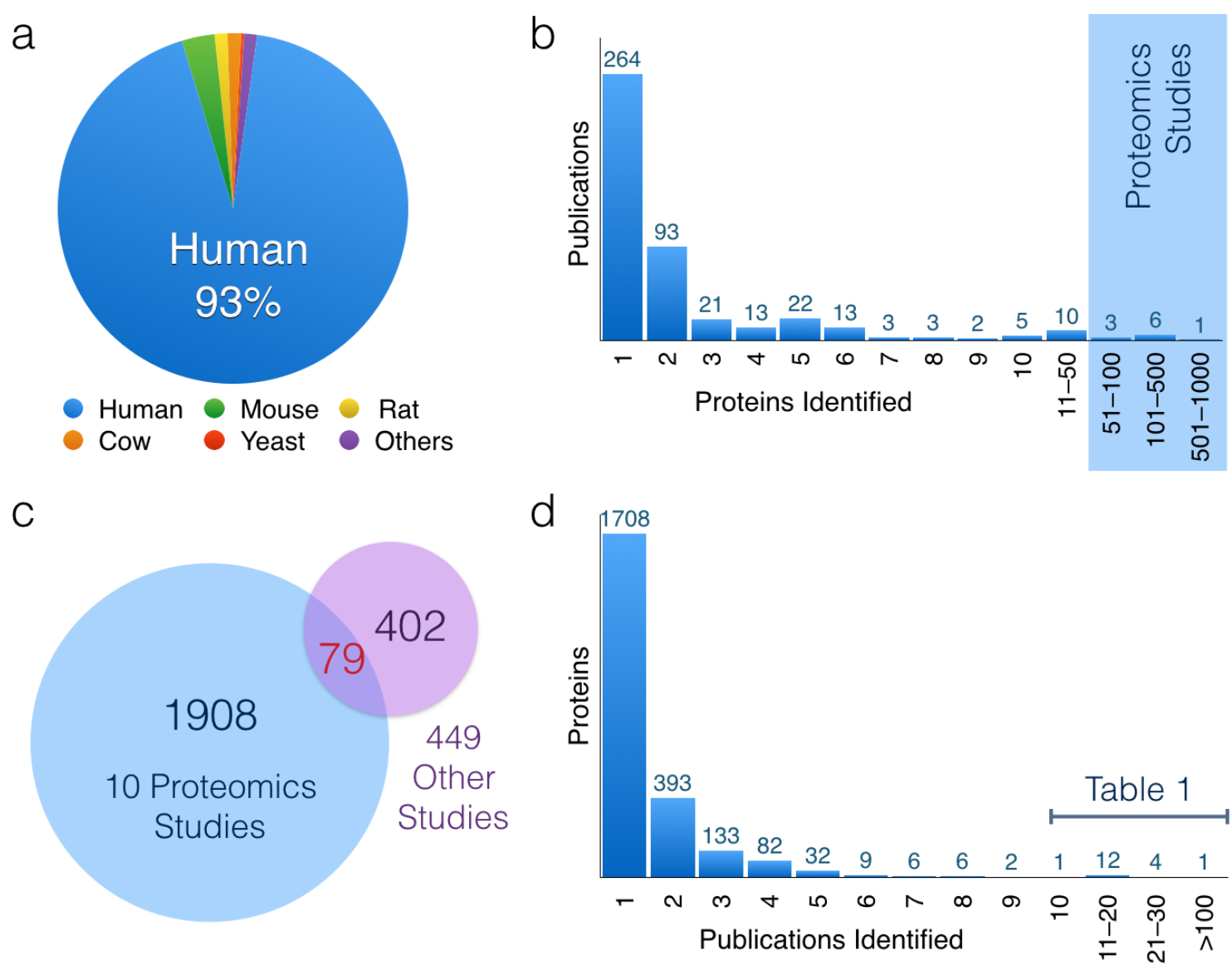
(a) The distribution of entries by species. (b) A histogram displaying how many modified proteins were identified per publication. The publications identifying the highest number of proteins are proteomics studies utilizing mass spectrometry. (c) Only 79 proteins were overlapped between 10 large mass spectrometry proteomics analyses (blue) and the remaining 449 studies (purple) included in the database. The relative lack of overlap indicates that current proteomics approaches have yet to reach saturation in identifying ADP-ribosylated substrates. (d) A representation of the number of times a specific protein is identified. The majority of proteins are only identified by one publication. However, 18 proteins were identified at least 10 times, and are listed in Table 1.

Amongst 459 publications, 58% of them (n=264) identified single ADP-ribosylated proteins, 5% (n=25) identified at least 10 substrates and 2% (n=10) from proteomics studies with more than 50 substrates identified (Figure 2b). Among the 2,389 unique proteins identified, 80% of ADP-ribosylated substrates (1,908) were uniquely identified in these 10 proteomics studies and 17% (402) from the rest of the 449 publications (Figure 2c). Surprisingly, only 79 ADP-ribosylated substrates were overlapped between these two sets, suggesting that we have yet to reach saturation in probing the ADP-ribosylated proteome. Consistently, >71% (1,708 out of 2,389 unique proteins) were identified only once (Figure 2d). ADP-ribosylation of human PARP1 (P09874) has been the most comprehensively analyzed protein with 111 publications, followed by histones, other PARPs and their homologs in mouse and rat (Table 1). Other proteins identified in at least 10 publications, including tumor suppressor p53 and DNA Topoisomerase 1, are also included in Table 1.

**Table 1.** A list of proteins identified independently in at least 10 publications. These 18 most identified proteins consist of a majority of PARPs and histones and their orthologues, along with DNA Topoisomerase 1 and the tumor suppressor p53.

| UniProt Entry | Protein Name | Number of Times Identified |
| --- | --- | --- |
| PARP1_HUMAN | Poly(ADP-ribose) polymerase 1 | 111 |
| H11_HUMAN | Histone H1.1 | 27 |
| H11_MOUSE | Histone H1.1 | 25 |
| PARP1_MOUSE | Poly(ADP-ribose) polymerase 1 | 24 |
| PARP1_RAT | Poly(ADP-ribose) polymerase 1 | 21 |
| P53_HUMAN | Tumor suppressor p53 | 18 |
| H31_HUMAN | Histone H3.1 | 18 |
| PARP1_BOVIN | Poly(ADP-ribose) polymerase 1 | 17 |
| H4_HUMAN | Histone H4 | 17 |
| H2A1_HUMAN | Histone H2A.1 | 16 |
| H2B1C_HUMAN | Histone H2B type 1-C/E/F/G/I | 15 |
| H11_RAT | Histone H1.1 | 14 |
| TNKS1_HUMAN | Tankyrase-1 | 13 |
| H10_HUMAN | Histone H1.0 | 12 |
| H2A2A_MOUSE | Histone H2A.2 | 12 |
| TOP1_HUMAN | DNA topoisomerase 1 | 11 |
| PARP2_HUMAN | Poly(ADP-ribose) polymerase 2 | 11 |
| H2B1A_RAT | Histone H2B type 1-A | 10 |

## ADP-ribosylation and non-membraneous RNA granules

The availability of this aggregate data potentially allows researchers to gain insights into global functions of ADP-ribosylation. Our previous informatics analyses indicated that ADP-ribosylated substrates are significantly enriched with proteins that are enriched for low-complexity regions, which aid self-assembly of non-membraneous organelles (25). Using 457 human proteins that have sequencing information in ADPriboDB, we tested whether these substrates are enriched in particular organelles. ADP-ribosylated substrates are significantly enriched in the proteome of non-membraneous RNA granules, including the nucleolus (38) and stress granules (39) (p-value = 1.85×10^−34^ and 5.00×10^−28^ respectively, Fisher’s Exact Test; Figure 3a). These data are consistent with the identification of the ADP-ribosyltransferases in these non-membraneous organelles–PARP1 and PARP2 in the nucleolus (40) and PARP5a, PARP12, PARP13, PARP14 and PARP15 in the stress granules (41). Consistent with a previous analysis (42), both the proteomes of the nucleolus and stress granules are enriched for low complexity domain-containing proteins (Figure 3a). In contrast, similar analyses for N-glycosylated substrates are enriched in the Endoplasmic Reticulum, Golgi, Lysosome and Membrane fractions, and these substrates are not enriched for low complexity domain-containing proteins (Figure 3a and b). Such a statistically significant association between ADP-ribosylated substrates and non-membraneous organelles is in agreement with our informatics-led hypothesis (25) and recent experimental evidence (43, 44), which indicate that PAR can help seed low complexity domain-containing proteins to form non-membraneous macromolecular structures.

**Figure 3.**
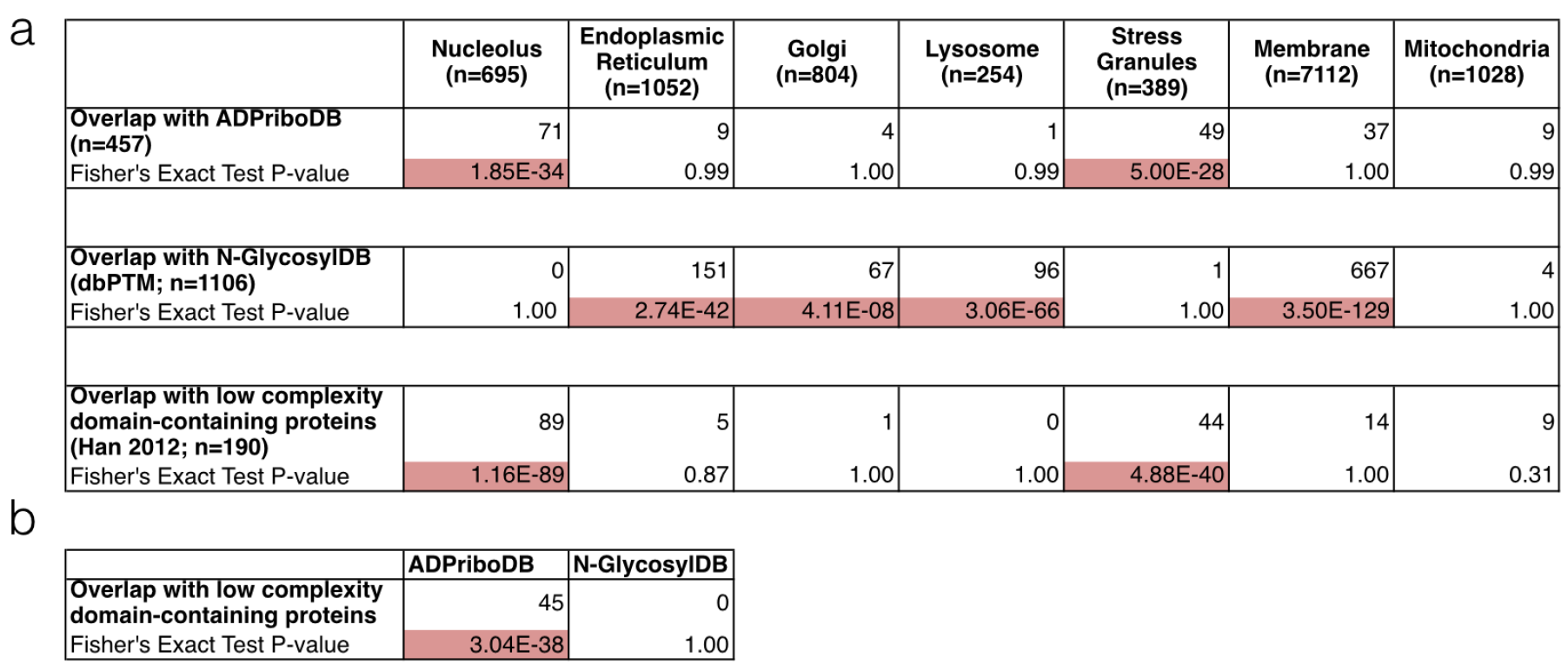
(a) ADP-ribosylated proteins are significantly enriched in non-membranous
compartments, such as the nucleolus (38) and stress granules (39), as determined by Fisher’s Exact Test, with significant p-values highlighted in red. These organelle proteomes are also enriched in low complexity domain-containing proteins. A similar analysis was performed with proteins that have experimentally validated N-glyosylation sites (N-GlyocsylDB) deposited in dbPTM (46). As expected, N-glyocsylated proteins were found enriched in the golgi, endoplasmic reticulum, lysosomes, and membrane fractions as defined by UniProtKB (51). No enrichment of experimentally validated ADP-ribosylation sites or N-glycosylation sites was found in the mitochondrial proteome (52). (b) Low complexity domain-containing proteins are significantly enriched in ADPriboDB in contrast to N-GlycosylDB, suggesting a link between ADP-ribosylation and low complexity domain-containing proteins.

## Future directions

ADPriboDB is built with the anticipation of several major developments in the ADP-ribosylation field–site identification, enzyme-substrate specificity, and substrate/site sensitivity to PARP inhibitors. First, given the development of various site identification methods (reviewed in (9, 14, 36)), we are expecting that a series of proteome-wide analyses will soon be available for a global survey of ADP-ribosylated proteins in cells and tissues. The first generation of ADPriboDB will therefore help connect these newly acquired proteomics data with the existing literature between 1975 and 2015. In addition, ADP-ribosylation sites can be used to correlate with existing databases on other post-translational modifications, such as phosphorylation, acetylation and methylation (45, 46). Second, 19% of ADPriboDB entries (2,419) include the information regarding the enzyme responsible for the modification of specific substrates, where 69% of these (1,659) are mediated by PARP1. However, we expect this trend will be changed soon with the coming-of-age technologies to investigate enzyme-substrate specificity of 17 human ADP-ribosyltransferases. These include proteome array (18, 19), the development of analog-sensitive ADP-ribosyltransferases (47–49) as well as the improvement in performing genetic knockdown by RNA interference and knockout using CRISPR in mammalian cell culture. Finally, PARP inhibition is actively used in clinical settings. Therefore, future development of these PARP-specific inhibitors for clinical use should couple with our basic scientific knowledge in how the endogenous ADP-ribosylated proteome is affected (9, 50). As a result, ADPriboDB starts to collect information under the field “Drug sensitivity” section to report whether certain substrates/sites are sensitive to particular PARP inhibitors (37). With ADPriboDB, we aim to provide a one-stop portal for the community to search ADP-ribosylation substrates and site information, where basic scientists and clinicians can use this cumulative information to understand basic biology of ADP-ribosylation, generate hypotheses in examining ADP-ribosylation functions, and explain clinical benefits and side effects observed in patients at the molecular level.

## ACKNOWLEDGEMENT

We thank Jerome Yu for the initial curation work, Diego Miranda-Saavedra for the discussion of ADPriboDB and the Leung lab members for critical discussions.

## Supplemental Information

**Table S1.** The standardized terms for the methods used in the 459 publications identifying ADP-ribosylated proteins in ADPriboDB. A diverse range of methods are utilized to identify modified proteins, ranging from chemical methods, such as the conjugation of NAD+ analogues, to more biochemical techniques, such as immunoprecipitation.

| Methods of characterizing ADP-ribosylation |
| --- |
| 2D Polyacrylamide Gel Electrophoresis (2D PAGE) |
| Autoradiography |
| Chromatography |
| Conjugation of 6-alkyne-NAD<sup>+</sup> |
| Conjugation of etheno-NAD<sup>+</sup> |
| Conjugation of mono-ADP-ribose to synthetic peptides |
| Conjugation of NAD<sup>+</sup> analogues |
| Dot blotting |
| Enzyme-linked immunosorbent assay (ELISA) |
| Fluorography |
| Immunofluorescence |
| In vitro assay |
| Immunoprecipitation (IP) |
| Mass Spectrometry |
| Microscopy |
| Mutagenesis |
| Polyacrylamide Gel Electrophoresis (PAGE) |
| Peptide Microarray |
| Phosphorimaging |
| Protein Microarray |
| Pulse Chase assay |
| Radioactivity assay |
| Slot blotting |
| Surface Plasmon Resonance |
| Thin Layer Chromatography (TLC) |
| Western blotting |

## REFERENCES

1. Hottiger,M.O. (2015) SnapShot: ADP-Ribosylation Signaling. Molecular Cell, 58, 1134–1134.e1.

2. Schreiber,V., Dantzer,F., Ame,J.-C. and de Murcia,G. de (2006) Poly(ADP-ribose): novel functions for an old molecule. Nat Rev Mol Cell Biol, 7, 517–528.

3. Vyas,S., Matic,I., Uchima,L., Rood,J.E., Zaja,R., Hay,R.T., Ahel,I. and Chang,P. (2014) Family-wide analysis of poly(ADP-ribose) polymerase activity. Nature Communications, 5, 4426.

4. Houtkooper,R.H., Pirinen,E. and Auwerx,J. (2012) Sirtuins as regulators of metabolism and healthspan. Nat Rev Mol Cell Biol, 13, 225–238.

5. Hottiger,M.O., Hassa,P.O., Lüscher,B., Schüler,H. and Koch-Nolte,F. (2010) Toward a unified nomenclature for mammalian ADP-ribosyltransferases. Trends Biochem Sci, 10.1016/j.tibs.2009.12.003.

6. Di Girolamo,M., Dani,N., Stilla,A. and Corda,D. (2005) Physiological relevance of the endogenous mono(ADP-ribosyl)ation of cellular proteins. FEBS J, 272, 4565–4575.

7. Cervantes-Laurean,D., Jacobson, E.L. and Jacobson, M.K. (1996) Glycation and glycoxidation of histones by ADP-ribose. J Biol Chem, 271, 10461–10469.

8. McDonald,L.J. and Moss,J. (1994) Enzymatic and nonenzymatic ADP-ribosylation of cysteine. Mol Cell Biochem, 138, 221–226.

9. Daniels,C.M., Ong,S.-E. and Leung,A.K.L. (2015) The Promise of Proteomics for the Study of ADP-Ribosylation. Molecular Cell, 58, 911–924.

10. Cervantes-Laurean,D., Jacobson,E.L. and Jacobson,M.K. (1997) Preparation of low molecular weight model conjugates for ADP-ribose linkages to protein. Methods in Enzymology, 280, 275–287.

11. Barkauskaite,E., Jankevicius,G. and Ahel,I. (2015) Structures and Mechanisms of Enzymes Employed in the Synthesis and Degradation of PARP-Dependent Protein ADP-Ribosylation. Molecular Cell, 58, 935–946.

12. Pascal,J.M. and Ellenberger,T. (2015) The rise and fall of poly(ADP-ribose): An enzymatic perspective. DNA Repair (Amst.), 10.1016/j.dnarep.2015.04.008.

13. Sharifi,R., Morra,R., Denise Appel,C., Tallis,M., Chioza,B., Jankevicius,G., Simpson,M.A., Matic,I., Ozkan,E., Golia,B., et al. (2013) Deficiency of terminal ADP-ribose protein glycohydrolase TARG1/C6orf130 in neurodegenerative disease. EMBO J, 10.1038/emboj.2013.51.

14. Vivelo,C.A. and Leung,A.K.L. (2015) Proteomics approaches to identify mono-(ADP-ribosyl)ated and poly(ADP-ribosyl)ated proteins. Proteomics, 15, 203–217.

15. Jungmichel,S., Rosenthal,F., Altmeyer,M., Lukas,J., Hottiger,M.O. and Nielsen,M.L. (2013) Proteome-wide Identification of Poly(ADP-Ribosyl)ation Targets in Different Genotoxic Stress Responses. Molecular Cell, 10.1016/j.molcel.2013.08.026.

16. Gagné,J.-P., Pic,E., Isabelle,M., Krietsch,J., Ethier,C., Paquet,E., Kelly,I., Boutin,M., Moon,K.-M., Foster,L.J., et al. (2012) Quantitative proteomics profiling of the poly(ADP-ribose)-related response to genotoxic stress. Nucleic Acids Res, 40, 7788–7805.

17. Dani,N., Stilla,A., Marchegiani,A., Tamburro,A., Till,S., Ladurner,A.G., Corda,D. and Di Girolamo,M. (2009) Combining affinity purification by ADP-ribose-binding macro domains with mass spectrometry to define the mammalian ADP-ribosyl proteome. Proc Natl Acad Sci USA, 106, 4243–4248.

18. Troiani,S., Lupi,R., Perego,R., Depaolini,S.R., Thieffine,S., Bosotti,R. and Rusconi,L. (2011) Identification of candidate substrates for poly(ADP-ribose) polymerase-2 (PARP2) in the absence of DNA damage using high-density protein microarrays. FEBS J, 278, 3676–3687.

19. Feijs,K.L., Kleine,H., Braczynski,A., Forst,A.H., Herzog,N., Verheugd,P., Linzen,U., Kremmer,E. and Lüscher,B. (2013) ARTD10 substrate identification on protein microarrays: regulation of GSK3β by mono-ADP-ribosylation. Cell Commun. Signal, 11, 5.

20. Gibson,B.A. and Kraus,W.L. (2012) New insights into the molecular and cellular functions of poly(ADP-ribose) and PARPs. Nat Rev Mol Cell Biol, 10.1038/nrm3376.

21. Feijs,K.L.H., Forst,A.H., Verheugd,P. and Lüscher,B. (2013) Macrodomain-containing proteins: regulating new intracellular functions of mono(ADP-ribosyl)ation. Nat Rev Mol Cell Biol, 14, 445–453.

22. Scarpa,E.S., Fabrizio,G. and Di Girolamo,M. (2013) A role of intracellular mono-ADP-ribosylation in cancer biology. FEBS J, 280, 3551–3562.

23. Hottiger,M.O. (2015) Nuclear ADP-Ribosylation and Its Role in Chromatin Plasticity, Cell Differentiation, and Epigenetics. Annu Rev Biochem, 10.1146/annurev-biochem-060614-034506.

24. Leung,A., Todorova,T., Ando,Y. and Chang,P. (2012) Poly(ADP-ribose) regulates post-transcriptional gene regulation in the cytoplasm. RNA Biol, 9.

25. Leung,A.K.L. (2014) Poly(ADP-ribose): An organizer of cellular architecture. The Journal of Cell Biology, 205, 613–619.

26. Teloni,F. and Altmeyer,M. (2016) Readers of poly(ADP-ribose): designed to be fit for purpose. Nucleic Acids Res, 44, 993–1006.

27. Krietsch,J., Rouleau,M., Pic,E., Ethier,C., Dawson,T.M., Dawson,V.L., Masson,J.-Y., Poirier,G.G. and Gagné,J.-P. (2012) Reprogramming cellular events by poly(ADP-ribose)-binding proteins. Mol. Aspects Med., 10.1016/j.mam.2012.12.005.

28. Perina,D., Mikoč,A., Ahel,J., Cetković,H., Zaja,R. and Ahel,I. (2014) Distribution of protein poly(ADP-ribosyl)ation systems across all domains of life. DNA Repair (Amst.), 10.1016/j.dnarep.2014.05.003.

29. Otto,H., Reche,P.A., Bazan,F., Dittmar,K., Haag,F. and Koch-Nolte,F. (2005) In silico characterization of the family of PARP-like poly(ADP-ribosyl)transferases (pARTs). BMC Genomics, 6, 139.

30. Rouleau,M., Patel,A., Hendzel,M.J., Kaufmann,S.H. and Poirier,G.G. (2010) PARP inhibition: PARP1 and beyond. Nat Rev Cancer, 10, 293–301.

31. Jagtap,P. and Szabó,C. (2005) Poly(ADP-ribose) polymerase and the therapeutic effects of its inhibitors. Nature reviews Drug discovery, 4, 421–440.

32. Feng,F.Y., de Bono,J.S., Rubin,M.A. and Knudsen,K.E. (2015) Chromatin to Clinic: The Molecular Rationale for PARP1 Inhibitor Function. Molecular Cell, 58, 925–934.

33. Lord,C.J., Tutt,A.N.J. and Ashworth, A. (2014) Synthetic Lethality and Cancer Therapy: Lessons Learned from the Development of PARP Inhibitors. Annu Rev Med, 10.1146/annurev-med-050913-022545.

34. Chambon,P., Weill,J.D. and Mandel,P. (1963) Nicotinamide mononucleotide activation of new DNA-dependent polyadenylic acid synthesizing nuclear enzyme. Biochem Biophys Res Commun, 11, 39–43.

35. Kraus,W.L. (2015) PARPs and ADP-Ribosylation: 50 Years … and Counting. Molecular Cell, 58, 902–910.

36. Rosenthal,F. and Hottiger,M.O. (2014) Identification of ADP-ribosylated peptides and ADP-ribose acceptor sites. Front Biosci, 19, 1041–1056.

37. Zhang,Y., Wang,J., Ding,M. and Yu,Y. (2013) Site-specific characterization of the Asp-and Glu-ADP-ribosylated proteome. Nat Methods, 10.1038/nmeth.2603.

38. Leung,A.K.L., Trinkle-Mulcahy,L., Lam,Y.W., Andersen,J.S., Mann,M. and Lamond,A.I. (2006) NOPdb: Nucleolar Proteome Database. Nucleic Acids Res, 34, D218–20.

39. Jain,S., Wheeler,J.R., Walters,R.W., Agrawal,A., Barsic,A. and Parker,R. (2016) ATPase-Modulated Stress Granules Contain a Diverse Proteome and Substructure. Cell, 164, 487–498.

40. Meder,V.S., Boeglin,M., de Murcia,G. and Schreiber,V. (2005) PARP-1 and PARP-2 interact with nucleophosmin/B23 and accumulate in transcriptionally active nucleoli. J Cell Sci, 118, 211–222.

41. Leung,A.K.L., Vyas,S., Rood,J.E., Bhutkar,A., Sharp,P.A. and Chang,P. (2011) Poly(ADP-ribose) regulates stress responses and microRNA activity in the cytoplasm. Molecular Cell, 42, 489–499.

42. Kato,M., Han,T.W., Xie,S., Shi,K., Du,X., Wu,L.C., Mirzaei,H., Goldsmith,E.J., Longgood,J., Pei,J., et al. (2012) Cell-free formation of RNA granules: low complexity sequence domains form dynamic fibers within hydrogels. Cell, 149, 753–767.

43. Altmeyer,M., Neelsen,K.J., Teloni,F., Pozdnyakova,I., Pellegrino,S., Grøfte,M., Rask,M.-B.D., Streicher,W., Jungmichel,S., Nielsen,M.L., et al. (2015) Liquid demixing of intrinsically disordered proteins is seeded by poly(ADP-ribose). Nature Communications, 6, 80–88.

44. Patel,A., Lee,H.O., Jawerth,L., Maharana,S., Jahnel,M., Hein,M.Y., Stoynov,S., Mahamid,J., Saha,S., Franzmann,T.M., et al. (2015) A Liquid-to-Solid Phase Transition of the ALS Protein FUS Accelerated by Disease Mutation. Cell, 162, 1066–1077.

45. Hornbeck,P.V., Zhang,B., Murray,B., Kornhauser,J.M., Latham,V. and Skrzypek,E. (2015) PhosphoSitePlus, 2014: mutations, PTMs and recalibrations. Nucleic Acids Res, 43, D512–20.

46. Huang,K.-Y., Su,M.-G., Kao,H.-J., Hsieh,Y.-C., Jhong,J.-H., Cheng,K.-H., Huang,H.-D. and Lee,T.-Y. (2016) dbPTM 2016: 10-year anniversary of a resource for post-translational modification of proteins. Nucleic Acids Res, 44, D435–46.

47. Carter-O’Connell,I., Jin,H., Morgan,R.K., Zaja,R., David,L.L., Ahel,I. and Cohen,M.S. (2016) Identifying Family-Member-Specific Targets of Mono-ARTDs by Using a Chemical Genetics Approach. Cell Rep, 14, 621–631.

48. Carter-O’Connell,I.O., Jin,H., Morgan,R.K., David,L.L. and Cohen,M.S. (2014) Engineering the substrate specificity of ADP-ribosyltransferases for identifying direct protein targets. J Am Chem Soc, 10.1021/ja412897a.

49. Gibson,B.A., Zhang,Y., Jiang,H., Hussey,K.M., Shrimp,J.H., Lin,H., Schwede,F., Yu,Y. and Kraus,W.L. (2016) Chemical genetic discovery of PARP targets reveals a role for PARP-1 in transcription elongation. Science, 10.1126/science.aaf7865.

50. Steffen,J.D., Brody,J.R., Armen,R.S. and Pascal,J.M. (2013) Structural Implications for Selective Targeting of PARPs. Front Oncol, 3, 301.

51. Boutet,E., Lieberherr,D., Tognolli,M., Schneider,M., Bansal,P., Bridge,A.J., Poux,S., Bougueleret,L. and Xenarios,I. (2016) UniProtKB/Swiss-Prot, the Manually Annotated Section of the UniProt KnowledgeBase: How to Use the Entry View. Methods Mol Biol, 1374, 23–54.

52. Smith, A.C. and Robinson, A. J. (2016) MitoMiner v3.1, an update on the mitochondrial proteomics database. Nucleic Acids Res, 44, D1258–61.

